# Identification of unknown proteins in X-ray crystallography and cryo-EM

**DOI:** 10.1101/2021.04.18.440303

**Authors:** Grzegorz Chojnowski, Adam J. Simpkin, Diego A. Leonardo, Wolfram Seifert-Davila, Dan E. Vivas-Ruiz, Ronan M. Keegan, Daniel J. Rigden

**Affiliations:** European Molecular Biology Laboratory, Hamburg Unit, Notkestrasse 85, 22607 Hamburg, Germany; Institute of Systems, Molecular and Integrative Biology, University of Liverpool, Liverpool L69 7ZB, England; São Carlos Institute of Physics, University of São Paulo, Avenida João Dagnone 1100, São Carlos, SP 13563-120, Brazil; European Molecular biology Laboratory. Meyerhofstraße 1, 69117, Heidelberg, Germany; Laboratorio de Biolgia Molecular, Facultad de Ciencias Biológicas, Universidad Nacional Mayor de San Marcos, Av. Venezuela Cdra 34 S/N, Ciudad Universitaria, Lima, Peru; UKRI-STFC, Rutherford Appleton Laboratory, Research Complex at Harwell, Didcot OX11 0FA, England

**Keywords:** protein structure, protein sequence, SIMBAD, crystallography, cryo-EM

## Abstract

Although experimental protein structure determination usually targets known proteins, chains of unknown sequence are often encountered. They can be purified from natural sources, appear as an unexpected fragment of a well characterized protein or as a contaminant. Regardless of the source of the problem, the unknown protein always requires tedious characterization. Here we present an automated pipeline for the identification of protein sequences from cryo-EM reconstructions and crystallographic data. We present the method’s application to characterize the crystal structure of an unknown protein purified from a snake venom. We also show that the approach can be successfully applied to the identification of protein sequences and validation of sequence assignments in cryo-EM protein structures.

## 1. Introduction

Recent years witnessed an unprecedented advancement of protein structure prediction approaches. Tools based on Deep Neural Networks proved not only much better than any previously available approaches, but also able to predict structures of proteins at a level of detail comparable to X-ray crystallography, which has been traditionally the predominant high-resolution technique (Jumper *et al*., 2020). Nevertheless, there are macromolecular targets which are not yet amenable to in-silico structure prediction approaches, most notably including structurally heterogeneous, large macromolecular complexes containing protein, RNA and small molecule components. Particularly interesting in this context are recent advances in cryogenic electron microscopy (cryo-EM) that enabled detailed studies of macromolecular complexes in their natural, cellular environment (Tegunov *et al*., 2021). The biochemical factors affecting the macromolecular composition and conformation are, however, not the only issue here as whole macromolecular complexes or their components may remain uncharacterized prior to structure determination attempts, or when purified from endogenous sources (Roh *et al*., 2018). The problem of unknown protein identity is not unique to cryo-EM studies. It is surprisingly common for macromolecular crystallographers to crystallize and solve previously uncharacterized protein structures. These can be proteins purified from natural sources (as described in this work) or contaminants, either native to an expression host (Niedzialkowska *et al*., 2016) or from elsewhere (Keegan *et al*., 2016; Hatti *et al*., 2016). Finally, samples can be simply mislabeled during crystallization or a synchrotron trip, which we have witnessed surprisingly often.

The experimental method of choice for protein characterization is sequencing of a sample (preparation or a crystal) using mass-spectrometry. In the case of cryo-EM, however, this may reduce the number of plausible sequence alternatives, but not solve the sequence assignment problem entirely (Ramrath *et al*., 2018). It is also not applicable to crystallography in cases where a crystal and sample solution are no longer available and cannot be reproduced.

Solving the crystal structure of a mis- or unidentified protein is a major difficulty. When reliable phase estimates can be obtained from anomalous differences, or directly from ultra-high-resolution diffraction data an initial model can be built based on an experimentally phased electron-density map. The use of a standard alternative phasing method - Molecular Replacement (MR) - is typically impossible due to lack of model selection criteria. In such a case a brute-force MR approach such as that implemented in SIMBAD (Simpkin *et al*., 2020, 2018), marathonMR (Hatti *et al*., 2017), or by the Wide-Search Molecular Replacement server (Stokes-Rees & Sliz, 2010) is the only option.

In a high-resolution crystal structure model the manual identification of protein sequences can be performed by an experienced crystallographer. However, similarly shaped amino-acid side-chains (e.g. Glutamate and Glutamine, or Aspartate and Asparagine) cannot usually be distinguished without an additional source of information. At lower resolutions the procedure becomes increasingly difficult, as model tracing itself is non-trivial without the sequence information (Chojnowski *et al*., 2019). In such cases, when at least a fragmented model can be traced in an electron-density map, the use of fold recognition tools (GESAMT (Krissinel, 2012), DALI (Holm & Laakso, 2016), or FatCat (Ye & Godzik, 2003)) may help to identify the protein (Niedzialkowska *et al*., 2016). This approach, however, may be difficult as the identification of complete protein chains in fragmented models, and in the presence of symmetry, is usually not possible without the use of target sequence. On the other hand, the use of short polypeptide stretches may be misleading as many remote-homologs tend to share structural motifs (Chojnowski *et al*., 2020).

For cryo-EM, the sequence identification methods mentioned above in the context of X-ray crystallography are generally applicable. Nevertheless, cryo-EM maps are usually determined at resolutions lower than in crystallography, making the model building and the identification of side-chain identities far more ambiguous and challenging (Chojnowski *et al*., 2021).There are, however, examples of successful protein identification that avoid these issues by a brute-force fitting of structures automatically predicted for a large pool of proteome sequences (Ramrath *et al*., 2018).

To the best of our knowledge there are two computer programs that can facilitate protein identification in X-ray crystallography and cryo-EM. Fitmunk (Porebski *et al*., 2016), originally side-chain modelling software, can assign probable residue identities to partial crystal-structure models to then query sequence databases using BLAST (Altschul *et al*., 1997). Although this approach generously assumes that the residue-type ambiguity can be modelled using standard scoring matrices hardcoded in BLAST, it was successfully used for the determination of sequence identity in several protein models (Niedzialkowska *et al*., 2016). CryoID (Ho *et al*., 2020), an approach designed to address the protein characterization in cryo-EM, uses the phenix.sequence_from_map (Terwilliger, 2003) tool to identify plausible residue identities using a six-letter sequence, simplified based on side-chain volume similarity. Similarly to Fitmunk, CryoID relies on standard sequence similarity scoring matrices that were adapted for the simplified six-letter sequence. It also requires manual curation and selection of most-reliable fragments, presumably due to the fact that by default the method strongly penalizes gaps in BLAST alignments and requires reliable, continuous main-chain traces on input. In a recent update (published when our paper was in submission) CryoID authors modified their protocol, which instead of BLAST can also use a more standard sequence alignment procedure that partially accounts for tracing errors in the models (Terwilliger *et al*., 2021).

It is known that tracing errors (deletions, insertions) are very common in intermediate models, in particular at lower resolutions and cryo-EM models, and correcting them usually requires the use of a target sequence (Chojnowski *et al*., 2019). Moreover, standard substitution matrices used for sequence alignment do not necessarily reflect the ambiguity of the residue types assigned based on partial models and maps. For example, in a recent study it was shown that a machine learning classifier most reliably discriminated shortest side-chains (alanine vs. glycine) that would fall into a single group simplified by side-chain size only (Chojnowski *et al*., 2019). To address these issues we developed *findMySequence*, a computer program that uses machine-learning predicted residue-type probabilities to query sequence databases using HMMER, a popular sequence analysis tool (Eddy, 2011). We show that the program successfully identifies sequences of protein models automatically built into crystallographic and cryo-EM maps, even though the automatically built models are usually highly fragmented and prone to tracing errors (Chojnowski *et al*., 2021). Furthermore, we use *findMySequence* to identify a protein purified and crystallized from a snake venom retrieved from *Bothrops atrox*, the most clinically important snake species in northern parts of South America (Estevao-Costa *et al*., 2016).

## 2. Materials and Methods

### 2.1. Crystal structures training set

For training the crystal structure residue-type classifier we selected protein crystal structures from the Protein Data Bank (PDB) (Berman *et al*., 2000). The selection criteria included pairwise sequence identity below 50% (PDB50 set), resolution between 2 and 3 Å, and crystallographic R-factor below 0.3. Out of 54,749 structures fulfilling these criteria on 4 February 2020 we selected 1,000 at random and downloaded corresponding “conservatively optimized” crystal structure models from the PDB_REDO server (Joosten *et al*., 2014) together with the corresponding map coefficients.

### 2.2. EM structures training set

For training the cryo-EM residue-type classifier we selected from PDB cryo-EM structures solved at a resolution better than 4 Å, with molecular weight below 500kDa, and half-maps available for download in EMDB (Velankar *et al*., 2016). As of 4 February 2020 we found 184 structures fulfilling these criteria. Initially, all the structures were refined into their corresponding maps with REFMAC5 (Murshudov *et al*., 2011*a*) using the auto_em.sh script from the ARP/wARP 8.1 suite (Chojnowski *et al*., 2021). Out of these, 117 models with CC_mask over 0.7 (estimated using phenix.map_model_cc,(Liebschner *et al*., 2019)) were selected for training the classifier.

### 2.3. Crystal structure identification benchmark set

For benchmarking the sequence identification procedure in crystal structures, we used a set of main-chain only models built using ARP/wARP for Molecular Replacement (MR) solutions at various target resolution and search-model similarity levels.

As targets we selected three hen egg-white lysozyme (HEWL) crystal structures solved over a range of resolutions: 2.9, 2.2 and 1.2 Å (PDB id codes 4gce, 4rln and 2hub respectively). All the target structures contain one molecule and 112 residues in the asymmetric unit and were solved in space group P43232, albeit with slightly different unit cell dimensions.

For each of the targets we used GESAMT (Krissinel, 2012) with default parameters to select a set of structurally similar models from the PDB (as of 4 February 2020). After excluding structures with CA-atoms only and those solved with powder diffraction or NMR, the sets contained 1 496, 1 478, and 1 492 models for targets at 2.9, 2.2, and 1.2 Å resolution respectively. The different number of search models can be attributed to small differences in the target structure coordinates. We used these as search models to solve the corresponding target structure with MR using Phaser (McCoy *et al*., 2007) with default parameters. The MR solutions were then used as an input for main-chain only model building with ARP/wARP 8.1. In addition to default model-building parameters, we employed a recently developed protocol for building short loops without sequence information (Chojnowski *et al*., 2019) to reduce model fragmentation.

### 2.4. EM structure identification benchmark set

To benchmark the sequence identification procedure in cryo-EM maps, we used two sets of ribosomal proteins; models built *de novo* and deposited models refined into corresponding EM maps.

From the PDB we selected cryo-EM structures of ribosomes determined at a resolution better than 3.5 Å with half-maps available for download in the Electron Microscopy Data Bank (EMDB). Initially, all the structures were refined into corresponding maps with REFMAC5 using the auto_em.sh script from ARP/wARP 8.1 suite (Chojnowski *et al*., 2021). Out of these we selected refined models with CC_mask over 0.7 (estimated using phenix.map_model_cc, (Liebschner *et al*., 2019)). The resulting set contained 17 ribosomes and 909 protein chain models originating from five different organisms: *Plasmodium falciparum, Escherichia coli, Staphylococcus aureus, Sus scrofa* and *Oryctolagus cuniculus*. For each of the protein chain models we estimated median local resolution for CA atom positions, using local resolution maps calculated using Resmap version 1.1.4 (Kucukelbir *et al*., 2014) with default parameters. Main-chain models for the test set proteins were traced fully automatically and *de novo* using ARP/wARP 8.1 with default parameters. To reduce model fragmentation we additionally used a recently developed protocol for building short loops without sequence information (Chojnowski *et al*., 2019). Each protein chain model was built in an artificial, rectangular box encapsulating a corresponding deposited model with a 5 Å margin. All remaining protein and RNA atoms from the corresponding ribosome model were masked with 3.0 Å radius.

### 2.5. Sequence databases

In principle, the protein sequence identification queries can be done against any set of sequences in FASTA format. Here, we used a set of 552,121 sequences corresponding to all protein chains available in PDB downloaded as of 17.09.2020 (PDB100). For the identification of ribosomal proteins in cryo-EM models we also used sets of reference proteomes downloaded from UniProt (The UniProt Consortium, 2021) for each of the targets. These were significantly smaller than the PDB100 set; three contained less than 6,000 sequences (*Plasmodium falciparum, Escherichia coli, Staphylococcus aureus*, UniProt ids UP000001450, UP000000625, and UP000008816 respectively), two mammalian proteomes had roughly 20,000 sequences each (*Sus scrofa, Oryctolagus cuniculus*, UniProt ids UP000008227 and UP000001811 respectively).

*Saccharomyces cerevisiae* proteome used for the identification of Voa1 assembly factor was downloaded from UniProt (UP000002311, 6,049 sequences). *Bothrops atrox* proteome sequences identified by Amazonas et al. (Amazonas *et al*., 2018) were downloaded from UniProt (selected by taxonomic identifier 8725).

### 2.6. Solving crystal structures with SIMBAD

In contrast to cryo-EM, where a complete map can be reconstructed from experimental data, the phases in crystallography are not measured and need to be retrieved from other sources (Fig. 1a). Molecular Replacement exploits the fact that evolutionarily related macromolecules tend to be structurally similar. Given sufficient similarity, a known structure correctly positioned in the target cell by MR can provide an approximation to the unknown phases of the target. In a typical MR search, suitable search models can be identified by a sequence search. However, in the case of the venom protein crystal discussed here, the target sequence was unknown. Therefore the domain database search option in *SIMBAD* was used to perform sequence-independent MR. This brute-force option makes use of the non-redundant domain database defined in the MoRDa application (Vagin & Lebedev, 2015) consisting of almost 100,000 domains from the PDB. These domains are used as search models in the rotation function step of MR as a quick means to score and identify possible homologues suited to providing the initial approximation to the target’s phases.

**Figure 1.**
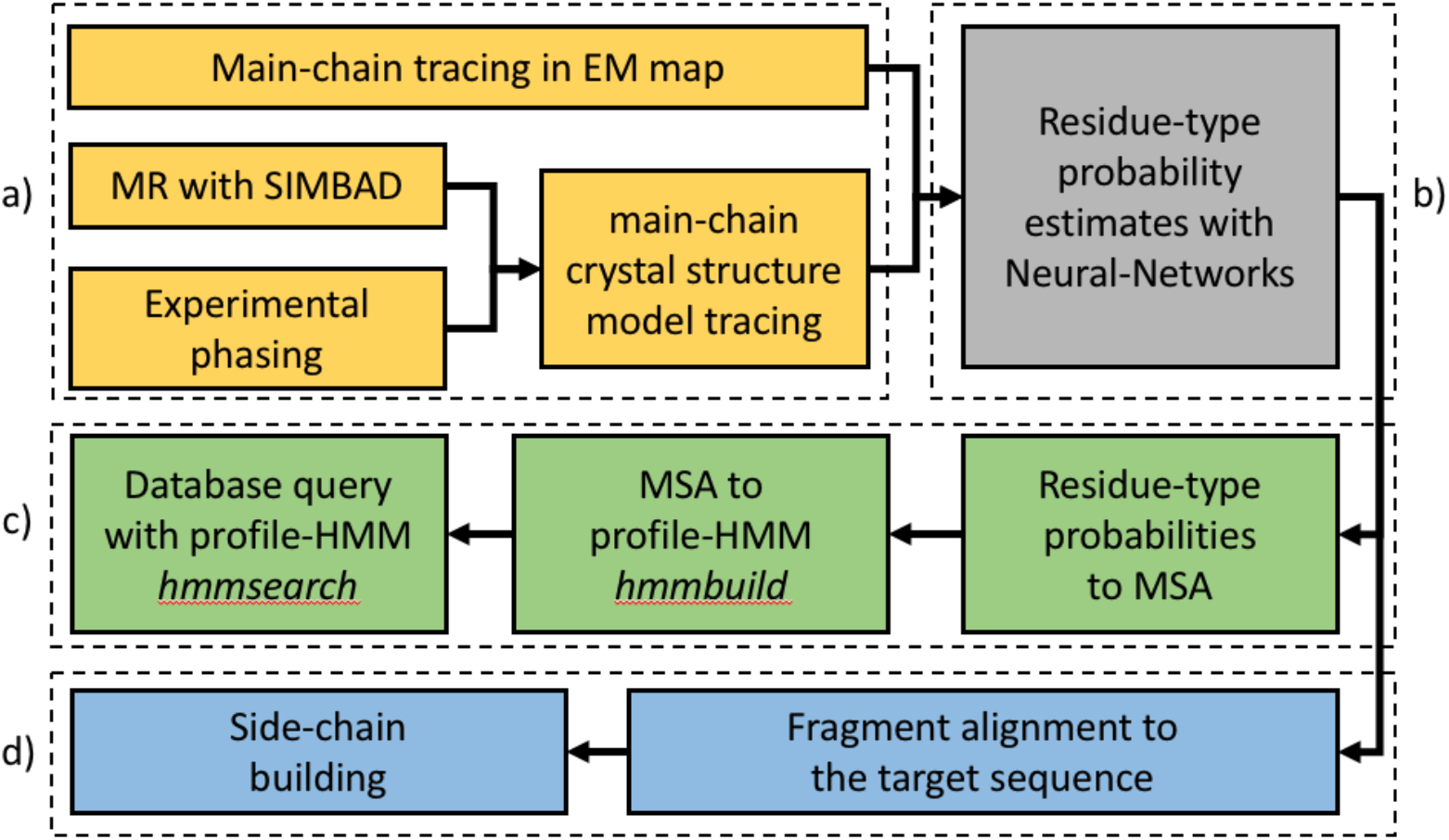
Schematic representation of the *findMySequene* usage workflow. Key steps are grouped in dashed rectangles; a) structure solution and model building, b) model interpretation, c) sequence database queries and d) sequence assignment and model building. All steps except model tracing (a) are integrated in the software and performed automatically.

### 2.7. Neural network model architecture and training

For predicting residue-type probabilities based on map values and main-chain models we built two neural-network models (Fig. 1b). The two models have identical architecture, but are trained on distinct training sets derived from crystal structures or cryo-EM models and their respective maps (see sections 2.1 and 2.2 for details).

Side-chain densities are described as a vector containing 324 map values sampled on a regular grid with 1.0 Å spacing (a residue descriptor). The grid is defined by N, CA, C peptide backbone atoms; centered at the CA atom and spanned by orthonormal vectors defined by N-CA atoms 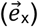, N-CA-C plane normal 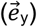, and their cross product 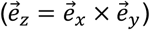. The input to the classifier contains all grid points that are within 1.0 Å distance from any side-chain atom in the top500 rotamers library (Lovell *et al*., 2000) aligned by N, CA, and C backbone atoms.

The neural-network model input is a vector of length 324 (the residue descriptor). The model contains two, fully connected hidden layers, the first with ReLU activation and 324 output features. The second layer has 20 output features and uses the log-softmax activation function enabling estimation of output classification probabilities. To avoid overfitting, we inserted an additional dropout layer with probability p=0.5 between the two hidden layers. The models were trained for 1,000 epochs with a batch size of 20, learning rate of 1e-4, and 10% validation set. For model training we used 617,477/68,608 and 177,831/19,758 training/test-set residue descriptors for X-ray and EM models respectively. The training/test set accuracies of the resulting models were 0.86/0.88 and 0.59/0.64 for crystal structure and cryo-EM models respectively.

### 2.8. Making queries in sequence databases with HMMER

To find a sequence in a database that matches the predicted residue-type probabilities, we use sequence comparison tools from the HMMER suite (Eddy, 2011)(Fig. 1c). Initially, predicted residue-type probabilities are converted into a multiple sequence alignment (MSA), where fractions of residue types in each column correspond to predicted probabilities. The residues in the input model are processed sequentially, starting from the longest continuous chain fragments. Next, the MSA is converted into a profile Hidden Markov Model (profile-HMM) using the default configuration of the *hmmbuild* program from the HMMER suite. Finally, the profile-HMM is used to query a sequence database using *hmmsearch* with default parameters and a sequence with the lowest, best-single-domain E-value is returned to the user.

### 2.9. Sequence assignment in main-chain models

To build side-chains in the input main-chain model fragment (Fig. 1c), we consider all possible alignments of a fragment to the target sequence. The most plausible alignment, given predicted residue-type probabilities, is then used to assign residue-types to the fragment. For computational efficiency, we consider only alignments of the fragments as a whole, which ignores tracing errors (insertions, deletions, or wrong connections).

Individual alignments are scored with a sum of log-probabilities of finding a specific residue type at consecutive positions in a chain fragment. Although our machine learning classifier has been calibrated and the predicted residue-type probabilities generally reflect expected frequencies, the accuracy of predictions may vary depending on resolution and quality of the models (Chojnowski *et al*., 2019). Therefore, for each alignment we calculate a standard score (Z-score) using log-probability distribution parameters estimated for a given fragment and a random target sequence. To additionally account for the varying target sequence length we estimate a probability that the highest Z-score selected from a number of alternative alignments of a fragment to the target sequence was observed by chance (p-value). For this purpose we apply Gumbel–Fisher–Tippett extreme-value distribution theorem using formulas derived previously for normalized structure factor amplitudes (Chojnowski & Bochtler, 2007). The analysis assumes that the Z-scores are normally distributed, which we confirmed on our benchmark set using the Shapiro-Wilk test at 99% confidence level (Shapiro & Wilk, 1965).

The extreme-value analysis allows for the comparison of scores obtained for multiple target sequences of various lengths. This, however, assumes that all the alternative alignments of a chain fragment to the target sequence are statistically independent, which is obviously not the case. To account for that, for p-value estimates we use a reduced number of alternative fragment alignments to the target sequence. According to our systematic analysis reducing this number by a factor of 10 results in p-values closest to the logistic-regression estimates of the correct assignment probabilities in our benchmark set.

### 2.10. Venom protein purification, crystallization, and data processing

A manual venom extraction on the main venom gland of *Bothrops atrox* from Pucallpa (Peru) was carried out. The venom was lyophilized and stored at -10 degrees until later use. A total of 350 mg lyophilized venom from *Bothrops atrox* was dissolved in 10 mL of 50 mM ammonium acetate buffer pH 5.0. The resuspended venom was centrifuged at 2,000g for 20 minutes at room temperature. The pellet containing insoluble particles was discarded. The clear supernatant was applied to a CM Sephadex ion exchange column C-50 (28 cm x 2.6 cm) previously equilibrated with the same buffer and the isocratic elution of unbound proteins was monitored at 280 nm. Bound proteins were eluted with a gradient (0 - 1M NaCl) at a flow rate of 1mL/min. In order to identify hemolytically active components, all eluted fractions were evaluated with the indirect hemolytic assay described previously (Camey *et al*., 2002). The fractions with the greatest activity, corresponding to the same peak, were pooled and concentrated for size exclusion purification. The concentrated fraction was applied to a Superdex 75 10/300 GL column, previously equilibrated with 50 mM ammonium acetate buffer pH 5.0. The enzymatic activity was monitored, and the purity of the protein was evaluated with SDS-PAGE.

The purified protein with hemolytic activity was concentrated to 11 mg/mL in 50 mM ammonium acetate buffer pH 5.0. Crystallization screening was performed with the sitting drop vapor diffusion method in 96-well plates using a Honeybee 931 robot (Genomic Solutions Inc.) and commercially available Crystal Screen II kit at 18 °C. After 7 days, single crystals (20% v/v 2-propanol, 20% w/v PEG 4000 and 0.1 M Sodium Citrate) were harvested and cryo-cooled in liquid nitrogen for data collection. X-ray diffraction data was collected on beamline MX-2 at the synchrotron radiation source at the Brazilian National Laboratory of Synchrotron Light from National Center for Energy and Materials (LNLS-CNPEM, Brazil) housing a PILATUS 2M detector. The data were indexed and integrated in iMosflm v.7.2.1 (Battye *et al*., 2011) and scaled with SCALA (Evans, 2006) to 1.95 Å resolution (Table 2).

### 2.11. Implementation and availability

The sequence identification program *findMySequence* was implemented using Python 3 with an extensive use of pytorch (Paszke *et al*., 2019), numpy (Oliphant, 2006), scipy (Virtanen *et al*., 2020), CCTBX (Grosse-Kunstleve *et al*., 2002) and CCP4 (Winn *et al*., 2011) libraries and utility programs. For making sequence database queries we use HMMER suite version 3.3.2. The program source code and installation instructions are available at https://gitlab.com/gchojnowski/findmysequence.

The model of the snake venom protein was deposited in Protein Data Bank with deposition code 7M6C.

## 3. Results

### 3.1. Benchmarks with deposited crystal structure models

To estimate the performance of our sequence identification procedure we used a large set of main-chain only protein crystal structure models built from MR solutions. We observed that the quality and completeness of a model is a main determinant of the success of the procedure. In cases when a reasonable model can be built (R-free below 50%) a largely correct sequence (above 80% target residues correctly assigned) can be identified in the vast majority of cases (Fig. 2a). Although this simple rule-of-thumb applies to all the tested targets, the performance of model building, and consequently the sequence identification, is clearly reduced at lower resolution (2.9 Å). This is in line with previous observations on the resolution dependence of the ARP/wARP model building results (Chojnowski *et al*., 2020).

**Figure 2:**
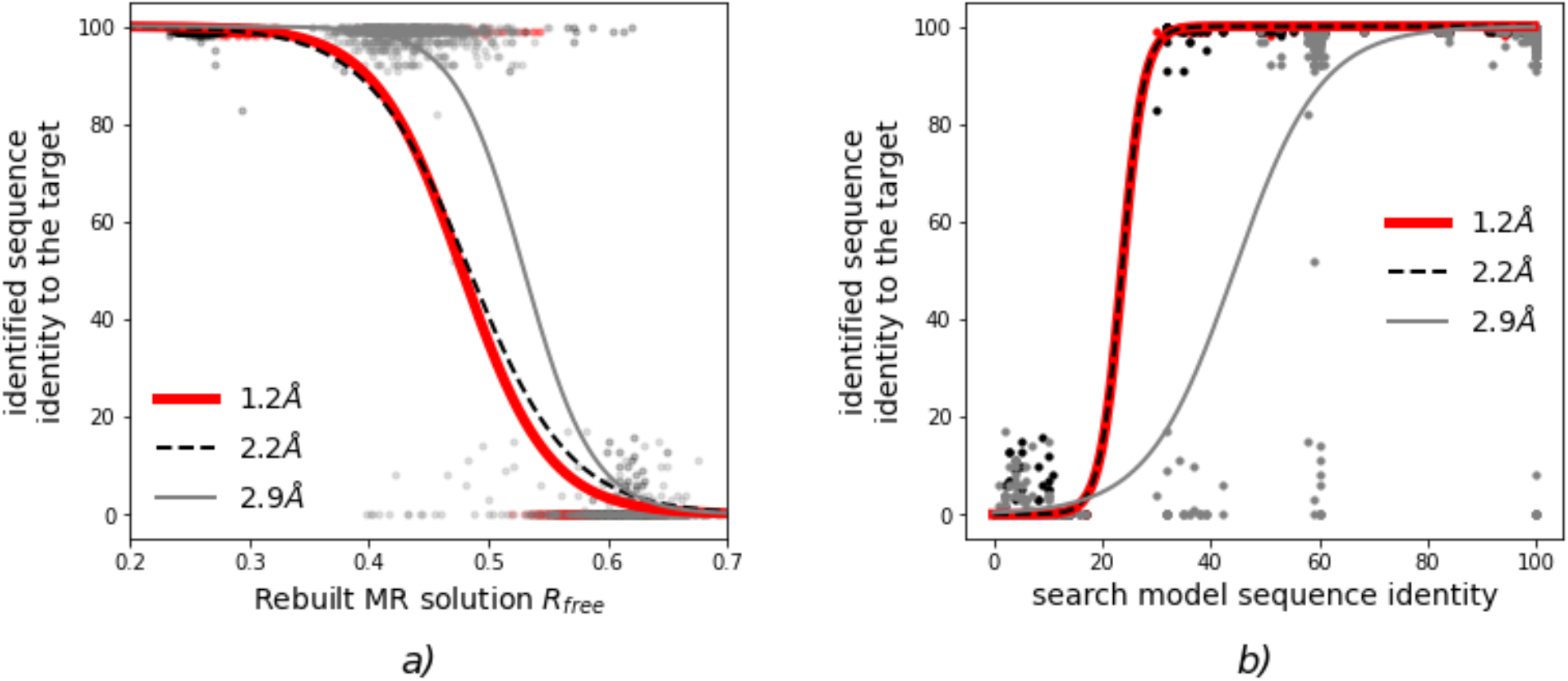
Sequence identification benchmarks for crystal structure models solved with MR using Phaser. Sequence identity of an identified sequence to the target sequence as a function of a) R-free factor value of a MR solution rebuild using ARP/wARP without input sequence and b) sequence identity of a MR search model to the target structure. The continuous and dashed curves are logistic-regression estimates of a probability that an identified sequence will have at least 80% sequence identity to the target sequence.

It must be noted that the R-free values observed here are higher than one would expect from a good quality model. This is due to the presence in the models of ‘free atoms’ used by ARP/wARP for the sparse representation of electron density maps. The atoms are not removed from final models built without sequence information (or with a low sequence coverage). As a consequence, complete main-chain only models built at lower resolutions are usually moderately overfitted, which results in high R-free values.

For all three targets a search model with at least 20% sequence identity is required to solve the structure and identify the corresponding sequence (Fig. 2b). However, for the lowest resolution dataset, sequence identification attempts with higher sequence-identity search models were also occasionally unsuccessful.

### 3.2. Benchmarks with cryo-EM ribosomal protein models

We benchmarked our automated sequence identification procedure on a set of 909 ribosomal proteins structures. We observed that a majority of sequences could be correctly identified using models built *de novo* up to 4.5 Å local resolution (Fig. 3a), which is usually too low for an automated method to trace a complete model (Lawson *et al*., 2021). The use of complete, deposited models readily increases sequence identification performance at lower local resolutions (Fig. 3b). We also observed that the sequence identification performance for models built *de novo* doesn’t depend on the size of a target sequence database (Fig. 3a).

**Figure 3:**
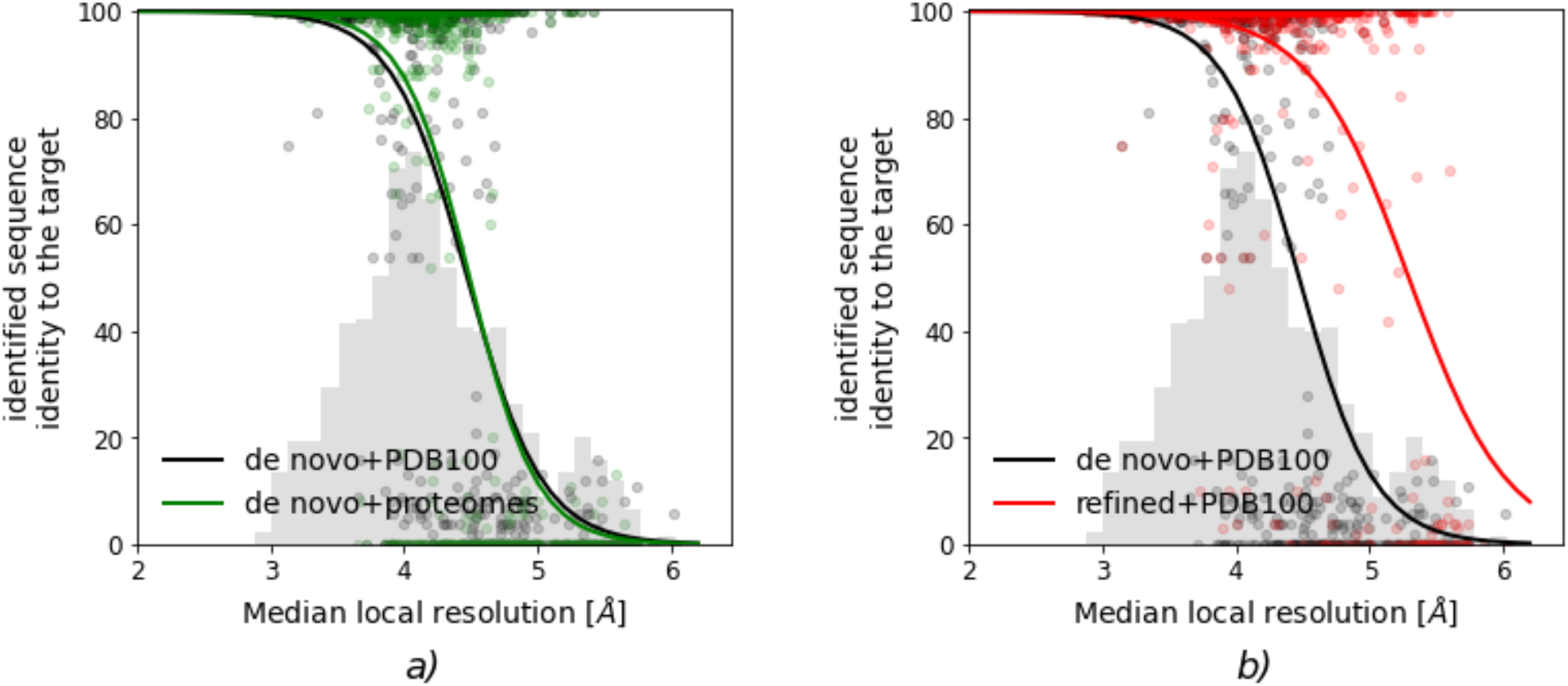
Sequence identification benchmarks for 909 cryo-EM models of ribosomal proteins. a) comparison of the method performances for an identification of models built *de novo* against small (proteomes) and large (PDB100) sequence databases b) comparison of the method performance for models built *de novo* and those based on refined deposited coordinates. Histograms of the median local resolution of the test set proteins are shown in grey (in arbitrary units). The continuous curves are logistic-regression estimates of a probability that an identified sequence will have at least 80% sequence identity to the target sequence.

We also evaluated a reliability of the best single domain E-value reported by HMMsearch (Fig. 4a). Generally, for E-values below 1e-7 a sequence close to the target was identified in most of the cases (logistic-regression estimate of correct identification probability exceeds 95%). At the same time, however, we observed that a number of hits with very low E-values do not match exactly the reference suggesting a limited, albeit very high, accuracy of the approach. In the case of crystal structure models, the logistic-regression estimate of the correct identification probability exceeds 95% for E-values below 1e-3. This is in line with our estimates of residue-type classifier accuracy, which is significantly higher for crystal than cryo-EM models.

**Figure 4:**
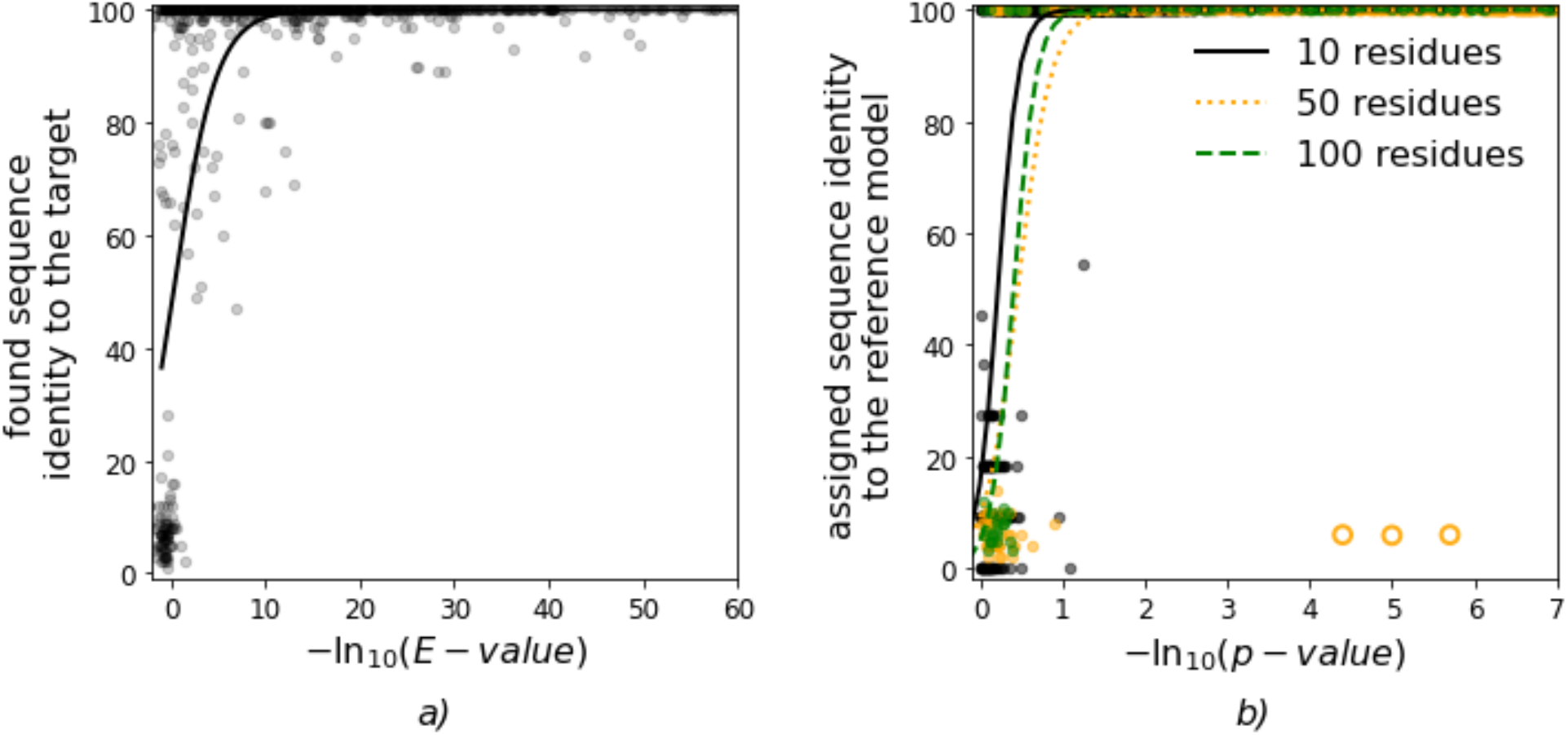
Sequence identification and assignment benchmarks for EM models. a) Identity of a sequence identified for models built *de novo* using ARP/wARP as a function of HMMsearch best-single-domain sequence alignment score b) Identity of a sequence assigned to continuous fragments of deposited EM models as a function of the sequence assignment score (p-value) for protein-fragment lengths of 10, 50 and 100 residues selected at random from test-set models. The continuous curves on the plots are logistic-regression estimates of a probability that an identified sequence will have at least 80% sequence identity to the reference model. Orange circles represent three reference chains with register error that were not used for the logistic-regression calculations.

### 3.3. Benchmarks of the sequence assignment in EM models

We validated the sequence assignment procedure using the complete test-set of ribosomal proteins. From the protein chain in the test set we selected random continuous fragments of lengths 10, 50 and 100 residues. For each fragment length we considered only sufficiently long protein chains, that resulted in 899, 820 and 548 fragments of length 10, 50 and 100 residues respectively. Next, we used our sequence assignment procedure to find an optimal alignment of the fragments to their deposited sequences given the corresponding map.

We observed that the p-value is a very accurate estimate of sequence assignment reliability and regardless of the model quality, local resolution, fragment and target sequence lengths for p-values below 1e-1 the sequence assignment is unambiguous (Fig. 4b).

In the sequence assignment results we encountered three clear outliers with p-value below 1e-4 and assigned sequences not matching the reference model (protein S21 in models of *E. coli* 70S ribosome at 3.0 Å resolution; pdb-id/emdb-id: 5we4/8814, 5wfs/8829, and 5wdt/8813). A closer look revealed that the reference models had obvious sequence register errors and therefore were removed for the benchmark set (Fig. 5). Interestingly, all three models were built based on an earlier model of *E. coli* 70S ribosome (pdb-id/emd-id: 5afi/2849 at 2.9 Å, not in our benchmark set) that also contains a register error in the corresponding chain.

**Figure 5.**
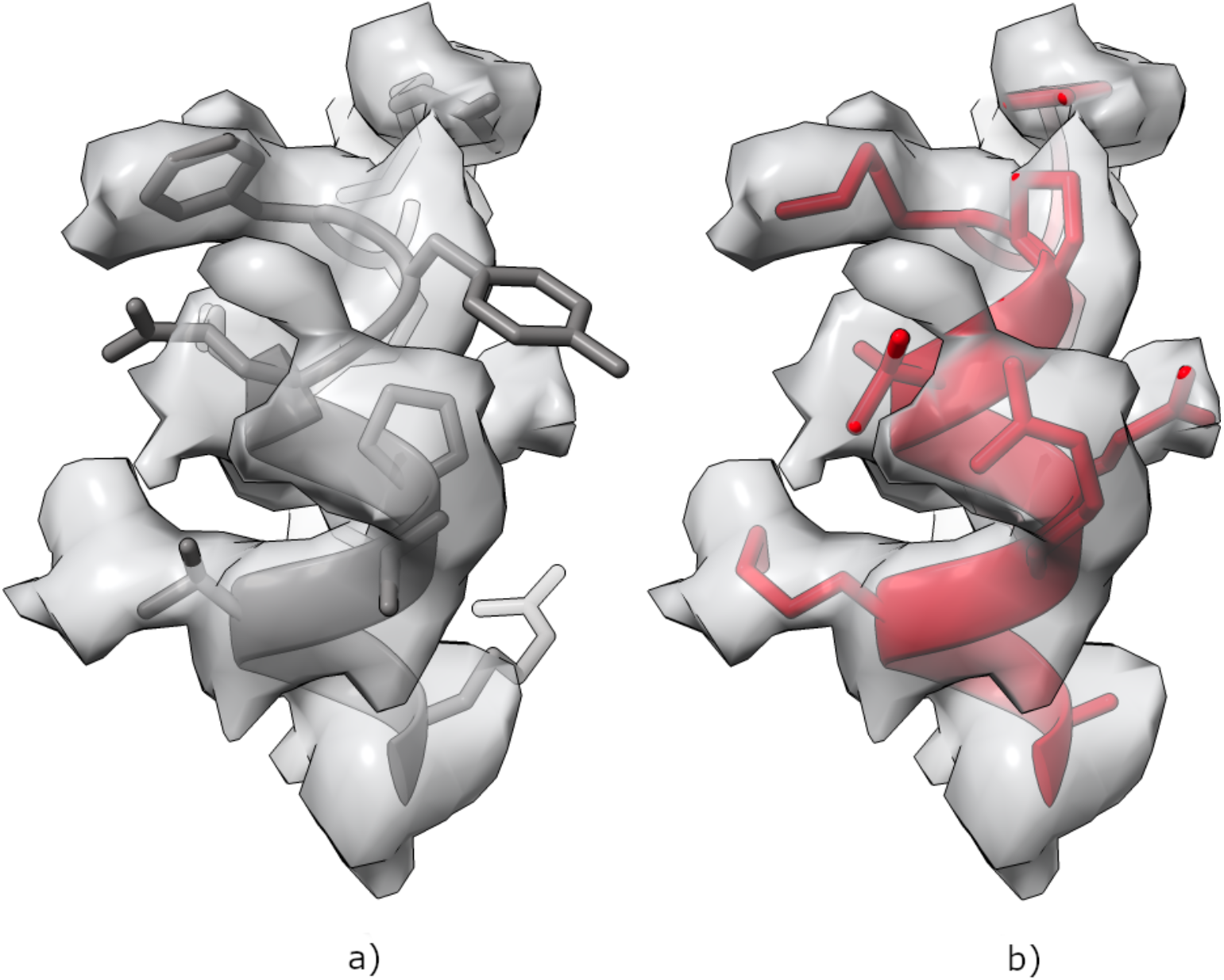
Fragment of a S21 protein model from *E. coli* 70S ribosome at 3.0 Å resolution (pdb-id/emdb-id: 5we4/8814). a) In the deposited coordinates, many side-chains outside a well-resolved map and a proline inside a regular alpha helix may raise suspicion a) After sequence re-assignment and side-chain rebuilding with *findMySequence* the map features are better explained by the model. Only residue range 34-44 in chain u and a corresponding map are shown for clarity.

### 3.4. Crystallography of proteins from natural sources

We collected diffraction datasets from crystal of a protein purified from *Bothrops atrox* that was observed to have phospholipase A2 (PLA2) activity. SIMBAD’s MoRDa database search was performed using default parameters and a model database created on September 18, 2019.

SIMBAD identified as a search model a PLA2 homologue purified from *Deinagkistrodon acutus* (pdb code: 1MC2), which agreed with the protein activity observed. *Phaser* (McCoy *et al*., 2007) placed one copy of the search model in the ASU with an LLG of 855 and a TFZ of 23.6. The model refined to R/R-free of 0.31/0.33 after 30 cycles of REFMAC5 (Murshudov *et al*., 2011*b*) refinement with jelly-body restraints. The moderate R/R-free factor values after refinement were related to differences between the search model and the target that were clearly visible in the map (Fig. 6a). An automated model rebuilding using ARP/wARP without input sequence improved main-chain traces in many regions (Fig. 6b). For the sequence identification we used a venom proteome determined previously for five specimens of *Bothrops atrox* snakes from two distinct Brazilian Amazon rainforest populations (Amazonas *et al*., 2018). Using *findMySequence* and an initial ARP/wARP model built without input sequence (Fig. 6b) we identified all six known PLA2 sequences in the venom proteome. The hmmsearch E-values for the hits clearly correlate with the sequence similarity of the sequences to the top-scored sequence variant (Table 1). A final model was built for the top scored sequence (A0A1L8D5Z7) that has 68% identity to the MR search model used to solve the structure. The crystal structure model was initially traced using ARP/wARP, rebuilt manually in COOT, and refined in REFMAC5 version 5.8.0267 using the CCP4 Cloud interface (Krissinel *et al*., 2018). The model refined to R/R-free of 0.19/0.23 (Fig. 6c). The data-collection and structure-refinement statistics are summarized in Table 2.

**Figure 6.**
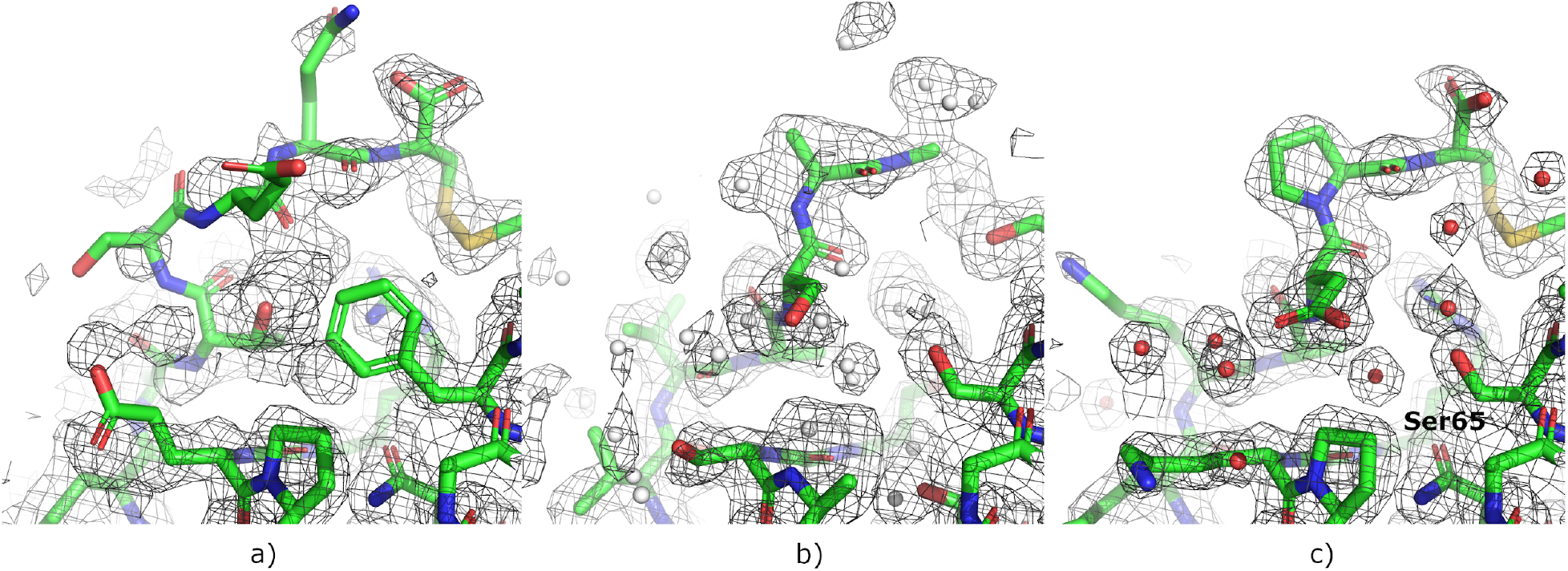
Consecutive steps of crystal structure determination and sequence identification of a protein with hemolytic activity purified from *Bothrops atrox* venom. a) Initial MR solution after 30 REFMAC5 refinement cycles with jelly-body restraints. The same fragment in b) ARP/wARP model re-traced without input sequence used as an input for *findMySequence* and c) final model. The 2Fo-Fc maps are contoured at the 1σ level above the mean. The free-atoms used for sparse electron-density map representation in ARP/wARP are shown as grey spheres. Water molecules are shown as red spheres.

**Table 1.**
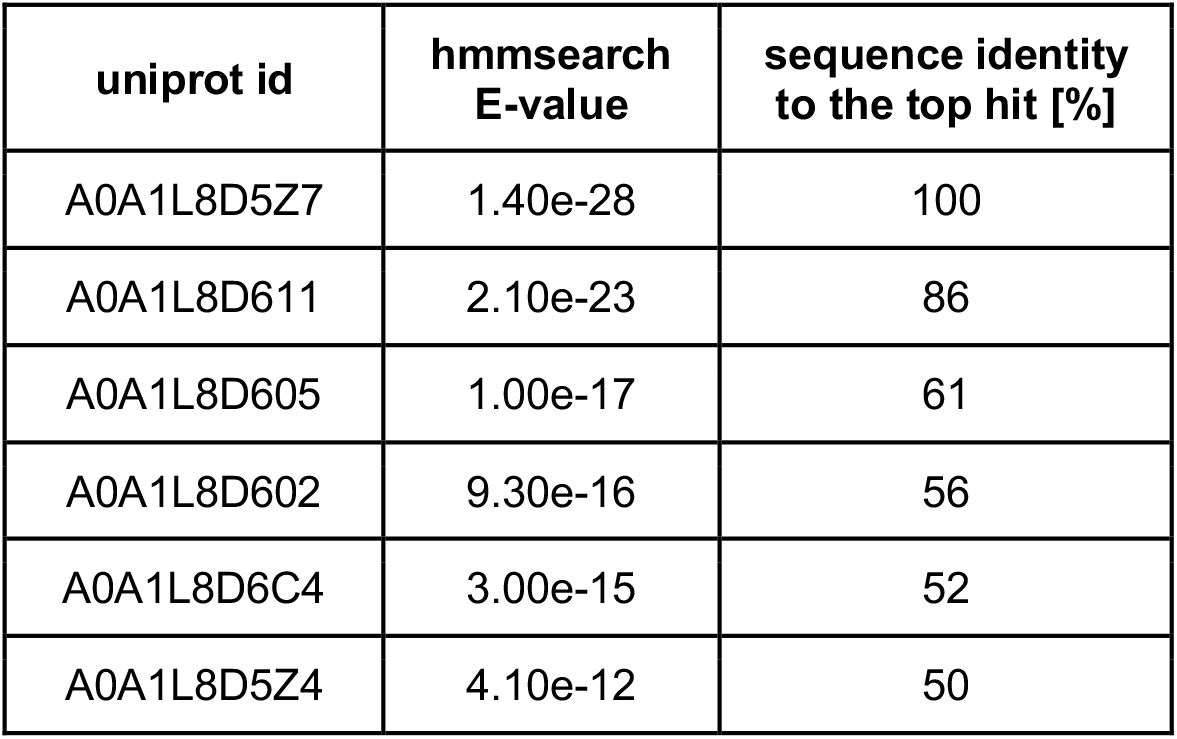
Results of the sequence identification in *Bothrops atrox* venom proteome using *findMySequence* and an ARP/wARP PLA2 model built without input sequence.

**Table 2.**
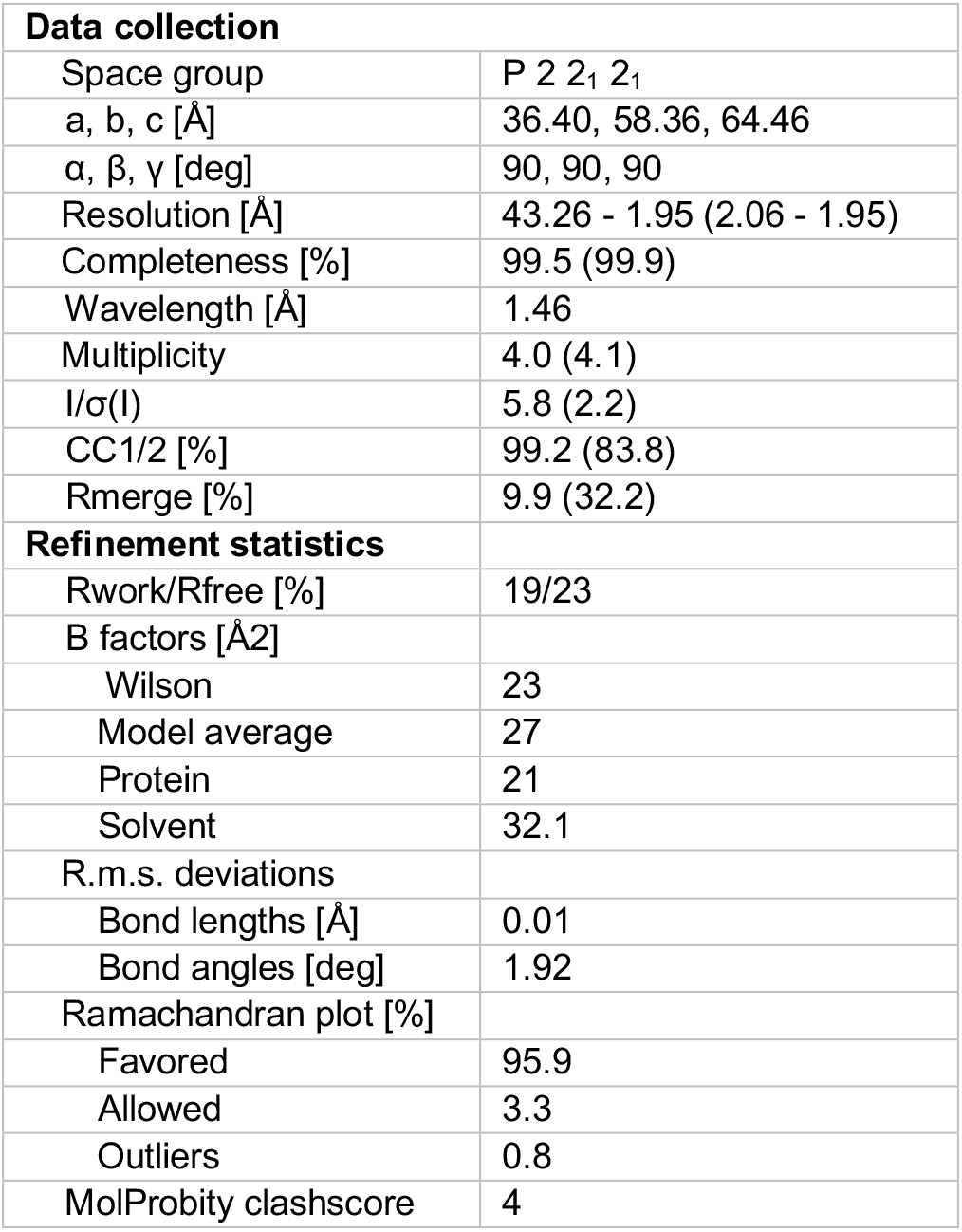
Data collection and structure refinement parameters for phospholipase A2 from *Bothrops atrox*. Lowest resolution shell values are given in brackets.

In the final, refined model we were able to identify a number of amino acids that unambiguously exclude the PLA2 sequence variants other than top scored A0A1L8D5Z7. These include a clearly resolved Ser65 (Fig. 6c), which in all other sequence variants is replaced with a Tyrosine or Phenylalanine. A careful, manual inspection of the final model did not reveal any possible sequence mismatches.

### 3.5. Identification of Voa1 assembly factor density in yeast V-ATPase Vo proton channel

A 3.5 Å cryo-EM structure of yeasts V-ATPase Vo proton channel revealed an α-helical density (Fig. 7) inside the central pore of the structure extending into the lumen of the cytoplasm suggesting that it may belong to a separate and unknown protein (Roh *et al*., 2018). The authors used mass spectrometry to identify this component as a Voa1 assembly factor. As deletion of the Voa1 was not required for the Vo assembly nor lethal, the result could be further confirmed with another EM reconstruction purified from a yeast strain with the Voa1 gene deleted, which was clearly missing the characteristic Voa1 density. We were able to unambiguously identify the Voa1 assembly factor with *findMySequence* in the yeast proteome (E-value 1.2e-13). We were also able to repeat the result with a single 27 residues long α-helix build into the Voa1 reconstruction using “place helix here” tool and real-space refined with secondary structure restraints in COOT (E-value of 2.4e-7 for correct and no result for reversed helix respectively). The result could be further confirmed with the sequence assignment procedure that unambiguously recognized the correct helix orientation and built the model side-chains (p-value 1.3e-3 and 0.9 for correct and reversed helix orientation respectively).

**Figure 7:**
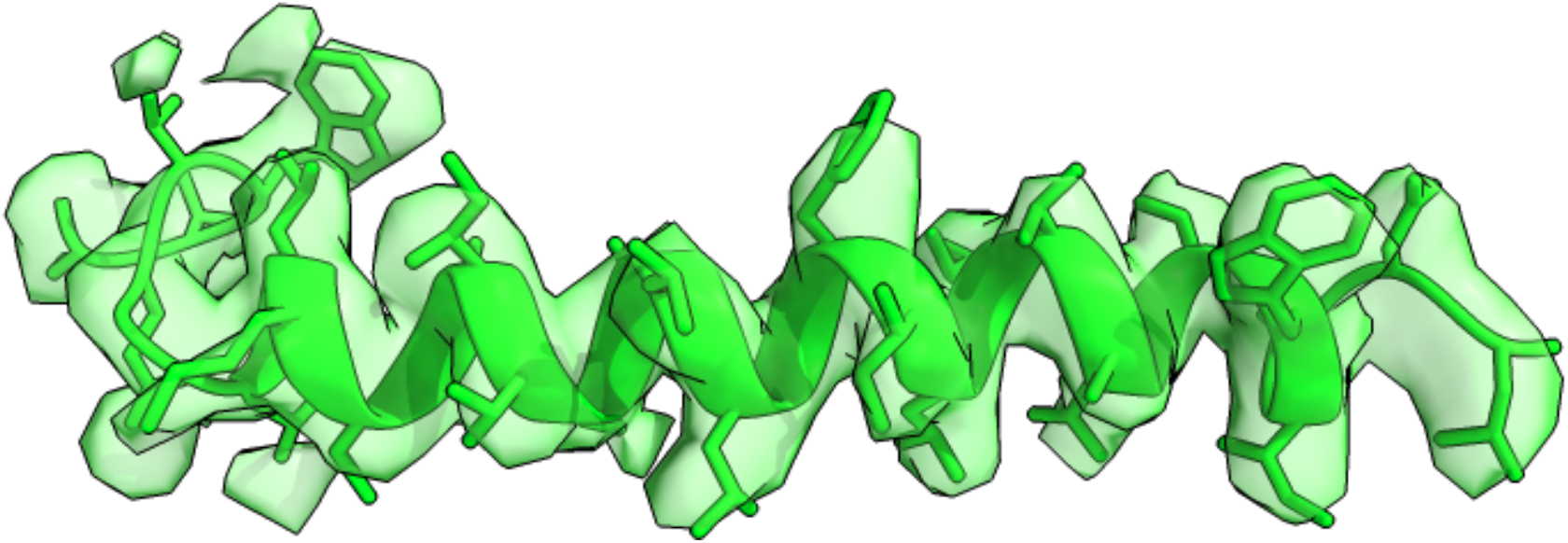
Model of Voa1 assembly factor and a corresponding cryo-EM reconstruction at 3.5 Å resolution (pdb and emdb codes 6c6l and 7348 respectively). Only a residue range 217-247 is shown for clarity.

### 3.6. Discussion and Conclusion

We have presented a complete pipeline for the identification of unknown proteins in crystallography and cryo-EM. A key element in the pipeline is a computer program *findMySequence* that identifies the most plausible protein sequence in a sequence database, given an electron-density map or cryo-EM reconstruction and a main-chain only model.

We show that our approach can successfully identify proteins, based on non-curated models automatically built into cryo-EM maps, at local resolutions up to 4.5 Å where models are usually highly fragmented and prone to tracing errors. We also show that the method’s performance increases for deposited coordinates and is then possible up to roughly 5.5 Å local resolution. In a more realistic, experimental setup, one can expect that the method performance will be comparable to the latter, as many errors in automatically traced cryo-EM models can be usually easily corrected manually. Therefore, we also expect that the use of model-building software other than ARP/wARP which we used in this work (e.g. phenix.map_to_model (Terwilliger *et al*., 2018), buccaneer (Hoh *et al*., 2020), MAINMAST (Terashi & Kihara, 2018), ROSETTA (Wang *et al*., 2016), or DeepTracer (Pfab *et al*., 2021)), should produce comparable results. It must be stressed, however, that most of the available programs require the target sequence on input and building a model before it becomes available may require an additional effort (e.g. the use of poly-ALA sequences). To address this issue we also show that alpha helices, which are relatively easy to build in medium resolution cryo-EM maps, are a good basis for the sequence identification procedure.

We observed that in cryo-EM structures our approach often identifies sequences that are very close, but not identical to the target. In crystallography, a sequence assignment can be validated after model refinement as residue-type mismatches usually produce elevated R-factor values and prominent difference density peaks. In cryo-EM, however, the sequence assignment validation is much more difficult. Although it often affects main-chain geometry, to the best of our knowledge, no automated tool exists that could be used for that purpose (Lawson *et al*., 2021). Therefore, we propose a sequence alignment procedure based on a machine-learning residue-type classifier used in this work. We have shown that it can unambiguously identify the correct sequence assignments of protein chains at a wide range of local resolutions. Even though at lower resolutions longer fragments may be needed to produce statistically relevant scores. We will describe this functionality of *findMySequence* in more detail in a separate publication. Nevertheless, it already proved crucial in building an atomic model of the mycobacterial ESX-5 type VII secretion system into a 3.4 Å resolution cryo-EM map (Beckham *et al*., 2020).

To further illustrate the practical use of *findMySequence* we describe the identification of a sequence of a protein purified and crystallized from a venom of a *Bothrops atrox* species snake. Prior to the crystallization, only the biochemical activity of the protein was known. Nevertheless, we were able to solve the structures using the brute-force MR approach implemented in SIMBAD and then identify the sequence using *findMySequence* with minimal manual input.

Our results show that availability of an initial model is a main limitation of the method application in crystallography. In our crystal-structure benchmark set every promising molecular replacement solution, with R-free below 50%, could be used for the successful identification of target sequences. This required a search model with at least 20% sequence identity to the target which agrees with a popular rule-of-thumb for search model suitability (Chojnowski *et al*., 2020). Thus, where a MR search is likely to be successful, the resulting map is likely to be tractable for *findMySequence*. Although at lower resolutions, this does not always guarantee success one may expect that owing to the number of available protein structures brute-force MR will often be able to provide an initial model for sequence identification (Simpkin *et al*., 2018). It should be noted that for benchmarks we used three good quality diffraction datasets, with data resolution being the major limiting factor. Crystallographers, however, often work with datasets bearing a number of pathologies like twinning or severe anisotropy. These, no doubt, would affect the performance of the approach and may require more effort from a crystallographer e.g. determination of experiential phases or manual model building. Nevertheless, the application of our method should save significant amounts of a structural biologist’s time in the majority of cases.

## 4. Acknowledgements

The development of the presented software was stimulated by students’ problems during numerous CCP4 and CCPEM workshops, where crystal structures and cryo-EM protein models often required characterization or challenging sequence assignment. We hope that the results obtained with the early prototypes of the software were as useful for them as they were for us.

We would like to thank Isabel Bento (EMBL) for valuable comments and critical reading of the manuscript and Florian K. M. Schur (IST Austria) for providing cryo-ET reconstructions used for testing the software.

This work was partially supported by the Biotechnology and Biological Sciences Research Council [BB/S007105/1].

## References

Altschul, S. F., Madden, T. L., Schäffer, A. A., Zhang, J., Zhang, Z., Miller, W. & Lipman, D. J. (1997). Nucleic Acids Res. 25, 3389–3402.

Amazonas, D. R., Portes-Junior, J. A., Nishiyama-Jr, M. Y., Nicolau, C. A., Chalkidis, H. M., Mourão, R. H. V, Grazziotin, F. G., Rokyta, D. R., Gibbs, H. L., Valente, R. H. & others (2018). J. Proteomics. 181, 60–72.

Battye, T. G. G., Kontogiannis, L., Johnson, O., Powell, H. R. & Leslie, A. G. W. (2011). Acta Crystallogr. Sect. D Biol. Crystallogr. 67, 271–281.

Beckham, K. S. H., Ritter, C., Chojnowski, G., Mullapudi, E., Rettel, M., Savitski, M. M., Mortensen, S. A., Kosinski, J. & Wilmanns, M. (2020). BioRxiv.

Berman, H. M., Westbrook, J., Feng, Z., Gilliland, G., Bhat, T. N., Weissig, H., Shindyalov, I. N. & Bourne, P. E. (2000). Nucleic Acids Res. 28, 235–242.

Camey, K. U., Velarde, D. T. & Sanchez, E. F. (2002). Toxicon. 40, 501–509.

Chojnowski, G. & Bochtler, M. (2007). Acta Crystallogr. Sect. A. 63, 297–305.

Chojnowski, G., Choudhury, K., Heuser, P., Sobolev, E., Pereira, J., Oezugurel, U. & Lamzin, V. S. (2020). Acta Crystallogr. Sect. D Struct. Biol. 76,.

Chojnowski, G., Pereira, J. & Lamzin, V. S. (2019). Acta Crystallogr. Sect. D Struct. Biol. 75, 753–763.

Chojnowski, G., Sobolev, E., Heuser, P. & Lamzin, V. S. (2021). Acta Crystallogr. Sect. D Struct. Biol. 77,.

Eddy, S. R. (2011). PLoS Comput. Biol. 7, e1002195.

Estevao-Costa, M. I., Gontijo, S. S., Correia, B. L., Yarleque, A., Vivas-Ruiz, D., Rodrigues, E., Chávez-Olortegui, C., Oliveira, L. S. & Sanchez, E. F. (2016). Toxicon. 122, 67–77.

Evans, P. (2006). Acta Crystallogr. Sect. D Biol. Crystallogr. 62, 72–82.

Grosse-Kunstleve, R. W., Sauter, N. K., Moriarty, N. W. & Adams, P. D. (2002). J. Appl. Crystallogr. 35, 126–136.

Hatti, K., Biswas, A., Chaudhary, S., Dadireddy, V., Sekar, K., Srinivasan, N. & Murthy, M. R. N. (2017). J. Struct. Biol. 197, 372–378.

Hatti, K., Gulati, A., Srinivasan, N. & Murthy, M. R. N. (2016). Acta Crystallogr. Sect. D Struct. Biol. 72, 1081–1089.

Ho, C.-M., Li, X., Lai, M., Terwilliger, T. C., Beck, J. R., Wohlschlegel, J., Goldberg, D. E., Fitzpatrick, A. W. P. & Zhou, Z. H. (2020). Nat. Methods. 17, 79–85.

Hoh, S. W., Burnley, T. & Cowtan, K. (2020). Acta Crystallogr. Sect. D Struct. Biol. 76, 531– 541.

Holm, L. & Laakso, L. M. (2016). Nucleic Acids Res. 44, W351–W355.

Joosten, R. P., Long, F., Murshudov, G. N. & Perrakis, A. (2014). IUCrJ. 1, 213–220.

Jumper, J., Evans, R., Pritzel, A., Green, T., Figurnov, M., Tunyasuvunakool, K., Ronneberger, O., Bates, R., Zidek, A., Bridgland, A. & others (2020). Fourteenth Crit. Assess. Tech. Protein Struct. Predict. (Abstract Book). 22, 24.

Keegan, R., Waterman, D. G., Hopper, D. J., Coates, L., Taylor, G., Guo, J., Coker, A. R., Erskine, P. T., Erskine, S. P. & Cooper, J. B. (2016). Acta Crystallogr. Sect. D Struct. Biol. 72, 933–943.

Krissinel, E. (2012). J. Mol. Biochem. 1, 76.

Krissinel, E., Uski, V., Lebedev, A., Winn, M. & Ballard, C. (2018). Acta Crystallogr. Sect. D Struct. Biol. 74, 143–151.

Kucukelbir, A., Sigworth, F. J. & Tagare, H. D. (2014). Nat. Methods. 11, 63–65.

Lawson, C. L., Kryshtafovych, A., Adams, P. D., Afonine, P. V, Baker, M. L., Barad, B. A., Bond, P., Burnley, T., Cao, R., Cheng, J. & others (2021). Nat. Methods. 18, 156–164.

Liebschner, D., Afonine, P. V, Baker, M. L., Bunkóczi, G., Chen, V. B., Croll, T. I., Hintze, B., Hung, L.-W., Jain, S., McCoy, A. J. & others (2019). Acta Crystallogr. Sect. D Struct. Biol. 75, 861–877.

Lovell, S. C., Word, J. M., Richardson, J. S. & Richardson, D. C. (2000). Proteins Struct. Funct. Genet. 40, 389–408.

McCoy, A. J., Grosse-Kunstleve, R. W., Adams, P. D., Winn, M. D., Storoni, L. C. & Read, R. J. (2007). J. Appl. Crystallogr. 40, 658–674.

Murshudov, G. N., Skubák, P., Lebedev, A. A., Pannu, N. S., Steiner, R. A., Nicholls, R. A., Winn, M. D., Long, F. & Vagin, A. A. (2011a). Acta Crystallogr. Sect. D Biol. Crystallogr. 67, 355–367.

Murshudov, G. N., Skubák, P., Lebedev, A. A., Pannu, N. S., Steiner, R. A., Nicholls, R. A., Winn, M. D., Long, F. & Vagin, A. A. (2011b). Acta Crystallogr. Sect. D Biol. Crystallogr. 67, 355–367.

Niedzialkowska, E., Gasiorowska, O., Handing, K. B., Majorek, K. A., Porebski, P. J., Shabalin, I. G., Zasadzinska, E., Cymborowski, M. & Minor, W. (2016). Protein Sci. 25, 720–733.

Oliphant, T. E. (2006). A guide to NumPy Trelgol Publishing USA.

Paszke, A., Gross, S., Massa, F., Lerer, A., Bradbury, J., Chanan, G., Killeen, T., Lin, Z., Gimelshein, N., Antiga, L. & others (2019). ArXiv Prepr. ArXiv1912.01703.

Pfab, J., Phan, N. M. & Si, D. (2021). Proc. Natl. Acad. Sci. 118,.

Porebski, P. J., Cymborowski, M., Pasenkiewicz-Gierula, M. & Minor, W. (2016). Acta Crystallogr. Sect. D Struct. Biol. 72, 266–280.

Ramrath, D. J. F., Niemann, M., Leibundgut, M., Bieri, P., Prange, C., Horn, E. K., Leitner, A., Boehringer, D., Schneider, A. & Ban, N. (2018). Science (80-.). 362,.

Roh, S.-H., Stam, N. J., Hryc, C. F., Couoh-Cardel, S., Pintilie, G., Chiu, W. & Wilkens, S. (2018). Mol. Cell. 69, 993–1004.

Shapiro, S. S. & Wilk, M. B. (1965). Biometrika. 52, 591–611.

Simpkin, A. J., Simkovic, F., Thomas, J. M. H., Savko, M., Lebedev, A., Uski, V., Ballard, C. C., Wojdyr, M., Shepard, W., Rigden, D. J. & others (2020). Acta Crystallogr. Sect. D Struct. Biol. 76, 1–8.

Simpkin, A. J., Simkovic, F., Thomas, J. M. H., Savko, M., Lebedev, A., Uski, V., Ballard, C., Wojdyr, M., Wu, R., Sanishvili, R. & others (2018). Acta Crystallogr. Sect. D Struct. Biol. 74, 595–605.

Stokes-Rees, I. & Sliz, P. (2010). Proc. Natl. Acad. Sci. 107, 21476–21481.

Tegunov, D., Xue, L., Dienemann, C., Cramer, P. & Mahamid, J. (2021). Nat. Methods. 18, 186–193.

Terashi, G. & Kihara, D. (2018). Nat. Commun. 9, 1618.

Terwilliger, T. C. (2003). Acta Crystallogr. - Sect. D Biol. Crystallogr. 59, 45–49.

Terwilliger, T. C., Adams, P. D., Afonine, P.V & Sobolev, O. V (2018). Nat. Methods. 15, 905.

Terwilliger, T. C., Sobolev, O. V, Afonine, P. V, Adams, P. D., Ho, C.-M., Li, X. & Zhou, Z. H. (2021). Acta Crystallogr. Sect. D Struct. Biol. 77, 1–6.

The UniProt Consortium (2021). Nucleic Acids Res. 49, D480–D489.

Vagin, A. & Lebedev, A. (2015). Acta Crystallogr. Sect. A Found. Adv. 71, s19–s19.

Velankar, S., van Ginkel, G., Alhroub, Y., Battle, G. M., Berrisford, J. M., Conroy, M. J., Dana, J. M., Gore, S. P., Gutmanas, A., Haslam, P. & others (2016). Nucleic Acids Res. 44, D385–D395.

Virtanen, P., Gommers, R., Oliphant, T. E., Haberland, M., Reddy, T., Cournapeau, D., Burovski, E., Peterson, P., Weckesser, W., Bright, J. & others (2020). Nat. Methods. 17, 261–272.

Wang, R. Y.-R., Song, Y., Barad, B. A., Cheng, Y., Fraser, J. S. & DiMaio, F. (2016). Elife. 5, e17219.

Winn, M. D., Ballard, C. C., Cowtan, K. D., Dodson, E. J., Emsley, P., Evans, P. R., Keegan, R. M., Krissinel, E. B., Leslie, A. G. W., McCoy, A., McNicholas, S. J., Murshudov, G. N., Pannu, N. S., Potterton, E. a, Powell, H. R., Read, R. J., Vagin, A. & Wilson, K. S. (2011). Acta Crystallogr. D. Biol. Crystallogr. 67, 235–242.

Ye, Y. & Godzik, A. (2003). Bioinformatics. 19, ii246.–ii255.

